# Ohana, a tool set for population genetic analyses of admixture components

**DOI:** 10.1101/071233

**Authors:** Jade Yu Cheng, Thomas Mailund, Rasmus Nielsen

## Abstract

**Motivation:** Structure methods are highly used population genetic methods for classifying individuals in a sample fractionally into discrete ancestry components.

**Contribution:** We introduce a new optimization algorithm of the classical Structure model in a maximum likelihood framework. Using analyses of real data we show that the new optimization algorithm finds higher likelihood values than the state-of-the-art method in the same computational time. We also present a new method for estimating population trees from ancestry components using a Gaussian approximation. Using coalescence simulations modeling populations evolving in a tree-like fashion, we explore the adequacy of the Structure model and the Gaussian assumption for identifying ancestry components correctly and for inferring the correct tree. In most cases, ancestry components are inferred correctly, although sample sizes and times since admixture can influence the inferences. Similarly, the popular Gaussian approximation tends to perform poorly when branch lengths are long, although the tree topology is correctly inferred in all scenarios explored. The new methods are implemented together with appropriate visualization tools in the computer package Ohana.

**Availability:** Ohana is publicly available at https://github.com/jade-cheng/ohana. Besides its source code and installation instructions, we also provide example workflows in the project wiki site.

## 1 Introduction

To quantify population structure, researchers often use methods based on the Structure model (Pritchard *et al.*, 2000). The basic assumption in this model is that individuals belong to a set of K discrete groups, each with unique allele frequencies and obeying Hardy-Weinberg Equilibrium, although the latter assumption can be relaxed (Gao *et al.*, 2007). Furthermore, individuals are allowed to have fractional memberships of each group. The groups are often termed ‘ancestry components’ and are sometimes interpreted to represent ancestral populations. This interpretation may be correct in some scenarios, for example when analyzing balanced samples of recently admixed individuals from otherwise highly divergent groups. However, if basic model assumptions are violated, for example if populations truly are not discrete units, the interpretation is more unclear. Nonetheless, inferences under the Structure model have proven highly popular for quantifying population genetic variation and for exploring the basic structure and divisions of genetic diversity in a sample.

STRUCTURE (Pritchard et al., 2000), FRAPPE (Tang et al., 2005), and ADMIXTURE (Alexander et al., 2009) are arguably the three most commonly used programs that apply the Structure model. STRUCTURE uses a Bayesian approach and relies on a Markov Chain Monte Carlo (MCMC) algorithm to sample jointly the posterior distribution of allele frequencies and fractional group memberships. FRAPPE uses a maximum likelihood approach and optimizes the likelihood for both allele frequencies and fractional group memberships using an expectation-maximization (EM) algorithm. ADMIXTURE uses the same model and statistical framework as FRAPPE but uses a faster optimization algorithm. ADMIXTURE executes a two-stage process, first taking a few fast EM steps and then executing a sequential quadratic programming (QP) algorithm. ADMIXTURE uses a pivoting algorithm to solve each QP problem and applies a quasi-Newton acceleration to each iteration. This acceleration does not respect parameter bounds. ADMIXTURE projects an illegal update to the nearest feasible point, and the acceleration step contributes only when it results in a better likelihood; otherwise the original QP update is used.

The interpretation of parameter estimates under the Structure model is somewhat contentious (Royal *et al.*, 2010;Weiss and Long, 2009). It is not clear exactly what the groups, or ancestry components, represent, but in the most simple interpretation we can think of them as estimates of some idealized ancestral populations. If a researcher has inferred the existence of K ancestral populations and knows the fractional memberships of each individual in these populations, a next question would be to explore their evolutionary history. The estimated allele frequencies can provide information about this.

The first approaches for using allele frequencies to estimate population histories dates back to the seminal work by Edwards and Cavalli-Sforza (Cavalli-Sforza *et al.*, 1964, 1967). They used Gaussian models for the joint distribution of allele frequencies of multiple populations to estimate genetic distances and to infer population trees. The use of Gaussian models to approximate genetic drift has recently had a resurgence after the availability of large Single Nucleotide Polymorphism (SNP) data sets. It is used in numerous methods and studies, including tests of local adaptation (e.g., (Coop *et al.*, 2010; Gunther *et al.*, 2013)) and the popular TREEMIX program developed by Pickrell *et al.* (2012). The basic idea in these methods is that you can define the joint allele frequencies among populations in terms of a Gaussian distribution with a covariance matrix dictated by a tree (or admixture graph). Under the Gaussian model, a tree corresponds to exactly one unique covariance matrix, and each covariance matrix corresponds to at most one tree. Furthermore, the likelihood function can be calculated very fast numerically without any need for pruning. The assumption of a Gaussian model for the allele frequencies corresponds to an assumption of a Brownian motion process to model genetic drift instead of, say, a Wright-Fisher diffusion. For small time intervals, the Brownian motion process can provide a close approximation to the Wright-Fisher diffusion. However, for longer time intervals, especially when the allele frequency is close to either of the boundaries (0 and 1), the Brownian motion model is clearly not a very accurate approximation to the Wright-Fisher diffusion. Nonetheless, the Gaussian models provide useful frameworks for inferences because of the distinct computational advantages.

A natural extension of the structure inference framework is to use similar models on the inferred ancestry groups to explore their evolutionary histories. A primary objective of this paper is to provide a computational tool for doing just this and to examine the performance of the Gaussian model in this context.

We present ‘Ohana’, a tool suite for inferring global ancestry, population covariances, and constructing population trees using Gaussian models. Ohana uses a maximum likelihood framework similar to ADMIXTURE, but it implements an optimization algorithm based on an Active Set (Murty *et al.*, 1988) method to solve the QP problem that, as we will show in the results section, tends to find higher maximum likelihood values than ADMIXTURE in similar computational time. In addition, using the model of NGSADMIX (Skotte *et al.*, 2013), it can work on genotype likelihoods from low coverage Next Generation Sequencing (NGS) data instead of called genotypes. It includes an optimization algorithm for estimating the best covariance matrix compatible with a tree, thereby estimating a tree, and simple algorithms and visualization tools for the obtaining a tree from the covariance matrix.

We evaluate the performance of the method on real and simulated data, and we also presents results on the limitations of the popular Gaussian model. We show, perhaps unsurprisingly, that the assumption of a Gaussian model in some cases can lead to severely biased branch lengths of population trees that have evolved under a Wright-Fisher diffusion process. This is a limitation of the approach implemented in Ohana and in other approaches that use Brownian motion models to approximate the Wright-Fisher diffusion.

## 2 Methods

Ohana’s **qpas** program infers admixture using genotype observations stored in the ped format from Plink (Purcell et al., 2007) or genotype likelihoods in the bgl format from beagle (Browning et al., 2007). Ohana’s **nemeco** program infers population covariances, and Ohana’s **convert** program facilitates different stages of the analysis by providing file conversions and fast approximations. The source code, installation instructions, and example workflows are available on GitHub at https://github.com/jade-cheng/ohana.

### 2.1 Statistical Models

The likelihood model using genotype observations is given by

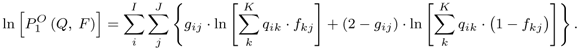

where *K* is the number of ancestry components, *I* is the number of individuals, and *J* is the number of polymorphic sites. This is the same as the model used in STRUCTURE (Pritchard *et al.*, 2000), FRAPPE (Tang *et al.*, 2005), ADMIXTURE(Alexander *et al.*, 2009), and SPA (Yang *et al.*, 2012).

Using the model in NGSADMIX (Skotte *et al.*, 2013), qpas can also work on genotype likelihoods. In that case the likelihood model is given by

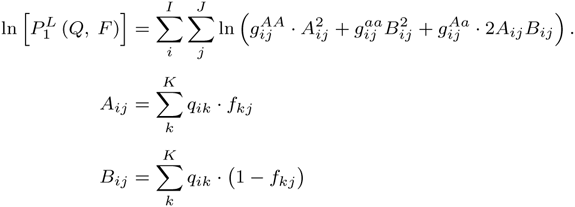

where 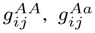, and 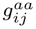 are the probabilities of observing the sequence data at the *i*th individual’s *j*th marker, conditioned on genotypes *AA*, *Aa* (or *aA*), and *aa*, respectively. This representation assumes markers with two alleles, although it could easily be generalized to multiple alleles. The advantage of working on genotype likelihoods instead of called genotypes is that genotype likelihoods incorporate the uncertainty regarding genotype calls inherent in much NGS data, and this makes it more applicable to low- or medium-coverage data (see e.g., (Skotte *et al.*, 2013)).

To infer population histories, Ohana models the joint distribution of allele frequencies across all ancestry components as a multivariate Gaussian similar to TREEMIX (Pickrell *et al.*, 2012) and Bayenv (Gunther *et al.*, 2013). The covariance matrix Ω of dimension *K* × *K* is assumed to be constant among all sites, and the process has a mean *μ_j_* at site *j*. The joint distribution of allele frequencies is then given by

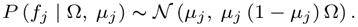

This system is under-determined (see e.g., (Felsenstein, 2004) chapter 23), i.e. multiple covariance matrices induce the same probability distribution on the allele frequencies. Similar to Felsenstein’s restricted maximum likelihood approach (Felsenstein, 1981), we therefore root the tree in one of the observations corresponding to conditioning on the allele frequencies in one of the populations when calculating the joint distribution of allele frequencies in the other populations. We emphasize that the rooting is arbitrary but that it does not imply any assumptions of this population actually being ancestral (for time reversible models). We then obtain a new covariance matrix Ω′, which has size (*K* – 1) × (*K* – 1) and a joint density of the form

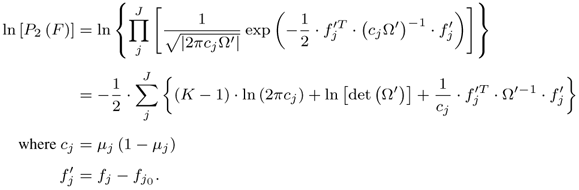

where *c_j_* = *µ_j_*(1–*µ_j_*)

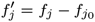.

### 2.2 Parameter Inference

#### 2.2.1 Inference for individual ancestries

To estimate *Q* and *F*, we use Newton’s approach. In general, we can approximate a function *F* (*x*) with its second order Taylor expansion. We proceed to minimize this second-order approximation by solving Δ*x*. In our problem, Δ*Q* and Δ*F* are constrained by 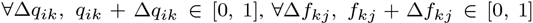, and 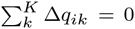 because 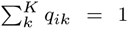. The analytical forms of the differential for 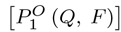 are presented below.

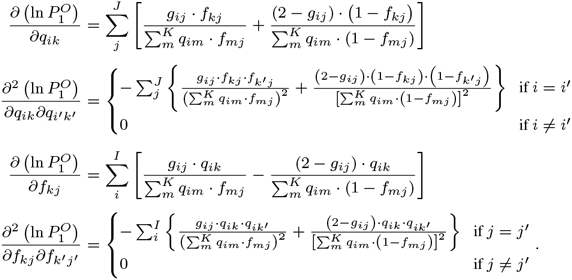

The analytical forms of the differential for 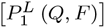 can also be found below. For both 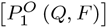 and 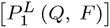, most off-diagonal values of the Hessians diminish. Leveraging this block structure, we convert the problem from manipulating huge matrices into manipulating sequences of small matrices of size *K*.

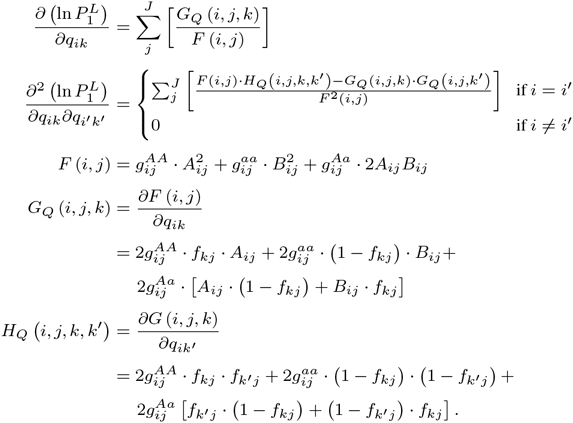

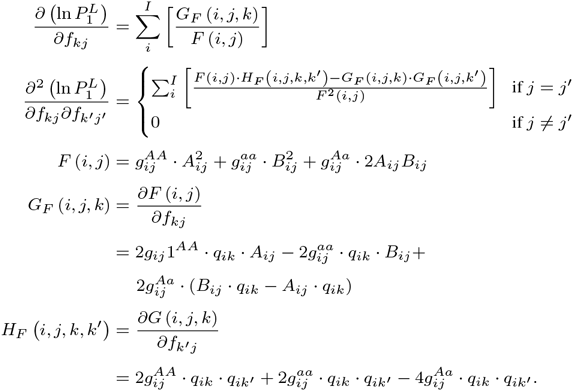

To solve these inequality- and equality-constrained quadratic optimization problems, we use an adaptation of the Active Set Algorithm (Murty *et al.*, 1988). To solve the equality problem defined by the active set and to compute the Lagrange multipliers of the active set, we use the Karush-Kuhn-Tucker (KKT) approach (Karush, 1939; Kuhn & Tucker, 1951). In each iteration, the algorithm searches for a better solution by considering the active constraints as equality constraints. It deviates from the bounds when the Lagrange multipliers signal a better solution toward the feasible region. The qpas program from Ohana performs this analysis. High-level pseudo-code of this algorithm appears in Algorithm 1 of the Supplementary Information (SI).

The maximum number of iterations performed by Ohana’s qpas to update *Q_i_* or *F_j_* is the number of constraints. In the worst case, the algorithm considers each constraint once. We have 2*K* + 1 constraints for updating *Q_i_* and 2*K* constraints for updating *F_j_*. Solving systems of linear equations used in KKT is at most Θ (*K*^3^). The runtime complexity for each update of *Q* and *F*, therefore, becomes 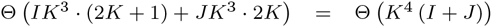, taking advantage of the block structure.

#### 2.2.2 Inference for population covariances

To optimize the likelihood model defined in the last equation of section 2.1, we use a black-box style of optimizer, the Nelder-Mead (NM) simplex method (Nelder & Mead *et al.*, 1965). We use sample covariances, 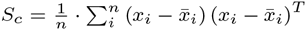, as the initial starting point for the NM optimizer, and we use Cholesky decomposition (Cholesky, 1910) to determine the positive semi-definiteness and to compute matrix inverses and determinants. The nemeco program in Ohana performs this analysis. High-level pseudo-code of this algorithm appears in SI Algorithm 2.

### 2.3 Estimation of phylogenetic trees

With the estimated covariance matrix in hand, we can construct a phylogenetic tree. We use the Neighbor-Joining (NJ) method for this, taking advantage of the NJ theorem (Saitou and Masatoshi, 1987), which states that when a distance matrix is compatible with a phylogentic tree, this tree will be accurately reconstructed by the NJ method. To do so, we first transform the covariance matrix to a distance matrix by observing the distance between two populations is given by Dist (*p*_1_, *p*_2_) = Var (*p*_1_) + Var (*p*_2_) – 2 × Cov (*p*_1_; *p*_2_).

Notice that there is a one-to-one correspondence between the covariance matrix and distances. These distances are then fed to the NJ algorithm. Ohana’s **convert** program performs all of these steps and in addition, provides an option to render the tree as SVG.

### 2.4 Simulated data

We used the software **fastsimcoal2** (Excoffier *et al.*, 2013) to produce genetic data using the Sequential Markov Coalescence (SMC) model (McVean and Niall, 2005; Marjoram and Simon, 2006). We simulated populations of nucleotide sequences according to a given demographic scenario. For each ancestry component, we simulated 100 sequences of size 20,000,000 bp under an identical population size of 50,000 for all components. We simulated demographic topologies with certain branch lengths by controlling population splits and effective population sizes.

We simulated admixture proportions for un-admixed and admixed scenarios. For un-admixed cases, we simply assigned a fraction of the sample to each population. For admixed cases, we simulated *Q_i_* independently from Dirichlet distributions Dir (*α*, *α*, *α*), similarly to the simulations used in (Pritchard *et al.*, 2000) and (Alexander *et al.*, 2009).

Finally, we also simulated genotype observations by first calculating the major allele frequency *f_ij_* for each individual at each marker location and then sampling genotypes under the assumption of Hardy-Weinberg Equilibrium, i.e. 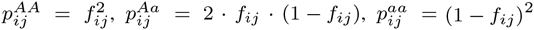, where 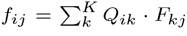, and *p^AA^*, *p^Aa^*, and *p^aa^* are the probabilities of observing major-major, major-minor, or minor-minor genotypes for the locus.

### 2.5 Real data

We used four data sets for the software comparison with ADMIXTURE shown in Figure 2 and Table 1:

- Dataset #1, a compilation of Europeans containing 17,507 markers and 118 individuals; this data was obtained from the POPRES (Nelson *et al.*, 2008), ALS (Laaksovirta *et al.*, 2010), Swedish Schizophrenia (Ripke *et al.*, 2013), and NCNG (Espeseth *et al.*, 2012) projects. It is a subset of data compiled for a study of Danish genetics
- Dataset #2, a compilation of HapMap (HapMap *et al.*, 2005) CEU, YRI, MEX, and ASW individuals containing 13,928 markers and 324 individuals. This is the benchmark dataset used in the original ADMIXTURE paper (Alexander *et al.*, 2009)
- Dataset #3, a compilation of Han Chinese samples from the HapMap project (HapMap *et al.*, 2005) containing 9,822 markers and 171 individuals.
- Dataset #4, a compilation of HapMap (HapMap *et al.*, 2005) world population of 4,695 markers 60 individuals of 10 North European, 10 Japanese, 10 Guaharati, 10 Luhya, 10 Maasai Kinyawa, and 10 Tuscan.

**Table 1.** A table of the highest log likelihoods achieved from ADMIXTURE and the qpas program in Ohana for a range *K* values. For each data set, each program, and each value of *K*, we executed 100 times using random seeds 0, 1,…, 99 and chose the highest value found in any run. This mimics the procedure often used for real data analysis. In the vast majority of cases, the qpas program in Ohana found significantly higher likelihood values than ADMIXTURE. Dataset #1 is a compilation of Europeans containing 17,507 markers and 118 individuals. Dataset #2 is the benchmark dataset used in ADMIXTURE (Alexander *et al.*, 2009) containing 324 CEU, YRI, MEX, and ASW individuals and 13,928 markers. Dataset #3 is a compilation of 171 Han Chinese samples and 9,822 markers. Dataset#4 is a worldwide population of 60 individuals and 4,695 markers.

| K | Dataset #1 |  |  | Dataset #2 |  |  | Dataset #3 |  |  | Dataset #4 |  |  |
| --- | --- | --- | --- | --- | --- | --- | --- | --- | --- | --- | --- | --- |
|  | Ohana | ADMIXTURE | Diff | Ohana | ADMIXTURE | Diff | Ohana | ADMIXTURE | Diff | Ohana | ADMIXTURE | Diff |
| 2 | -1967733 | -1967733 | 0 | -3835358 | -3835365 | 7 | -1857263 | -1857263 | 0 | -288991 | -288991 | 0 |
| 3 | -1956785 | -1956799 | 14 | -3799873 | -3799887 | 14 | -1848450 | -1848451 | 1 | -279462 | -279463 | 1 |
| 4 | -1946218 | -1946244 | 26 | -3788598 | -3788607 | 10 | -1841198 | -1841199 | 1 | -275212 | -275213 | 1 |
| 5 | -1935775 | -1936025 | 250 | -3777351 | -3777361 | 11 | -1834377 | -1834378 | 1 | -271807 | -271808 | 1 |
| 6 | -1925636 | -1925877 | 241 | -3766558 | -3766540 | -18 | -1827829 | -1827830 | 2 | -268837 | -268832 | -5 |
| 7 | -1915552 | -1915743 | 191 | -3755851 | -3755860 | 9 | -1821445 | -1821458 | 13 | -265907 | -265923 | 17 |
| 8 | -1905430 | -1905638 | 209 | -3746227 | -3745412 | -815 | -1815214 | -1815214 | 0 | -263052 | -263096 | 44 |
| 9 | -1895372 | -1895879 | 507 | -3735240 | -3736079 | 839 | -1809084 | -1809101 | 18 | -260268 | -260440 | 172 |
| 10 | -1885306 | -1885466 | 160 | -3725558 | -3725624 | 66 | -1802911 | -1802906 | -5 | -257539 | -257736 | 197 |
| 11 | -1875503 | -1875853 | 350 | -3715543 | -3715157 | -385 | -1796763 | -1796847 | 84 | -254920 | -254961 | 41 |
| 12 | -1865492 | -1865965 | 474 | -3706069 | -3707715 | 1646 | -1790671 | -1790811 | 140 | -252196 | -252266 | 70 |
| 13 | -1855502 | -1856262 | 760 | -3697531 | -3698519 | 987 | -1784688 | -1784765 | 77 | -249456 | -249468 | 12 |
| 14 | -1845732 | -1846490 | 758 | -3688970 | -3689124 | 154 | -1778599 | -1778671 | 73 | -246760 | -246817 | 56 |
| 15 | -1836315 | -1836775 | 460 | -3681092 | -3680829 | -263 | -1772555 | -1772669 | 114 | -244058 | -244298 | 240 |

For the admixture and covariance data analysis shown in Figure 5, we used a combination of world-wide samples containing 127,855 markers and 80 individuals from the HGDP project. We pruned for minor allele frequencies and Linkage Disequilibrium (LD) with Plink (Purcell *et al.*, 2007) using the options –*indep 50 5 2 –geno 0.0 –maf 0.05*.

## 3 Results

### 3.1 Computational speed

ADMIXTURE has previously been shown to have the most efficient optimization algorithm among the previously published methods (Alexander *et al.*, 2009). We therefore compare the optimization algorithm in Ohana to the algorithms implemented in ADMIXTURE. For a fair comparison, we show the distribution of likelihood values for the two methods, obtained after a fixed amount of computational time, for multiple different runs of Ohana and ADMIXTURE (Figure 2 and Table 1). We verify that the likelihood values are comparable between the two programs by calculating likelihood values for the same parameter values for both programs. We use four different real data sets described in the Methods section and explore a range of different values of *K*. For a very short amount of computational time, ADMIXTURE tends to find higher likelihood values. ADMIXTURE may possibly use better initial values for the optimization. However, after a relative short amount of time, the qpas algorithm in Ohana tends to find higher likelihood values than ADMIXTURE for the same computational time.

**Fig. 1.**
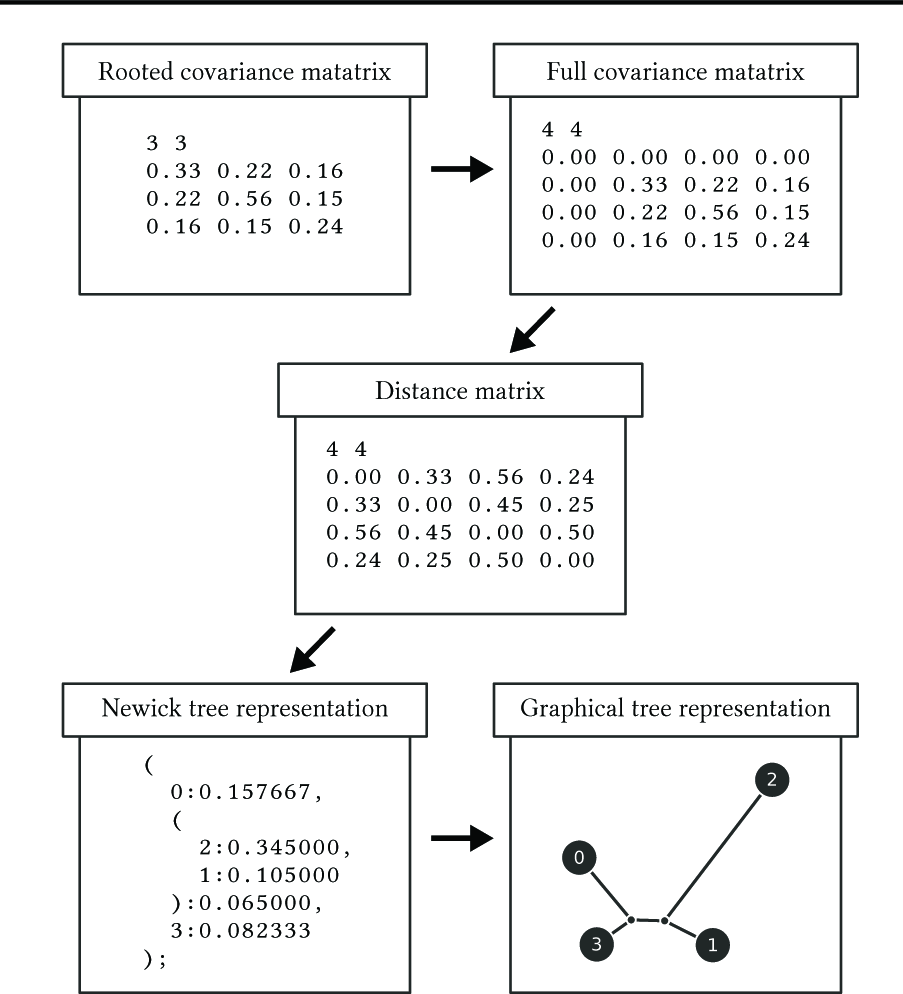
Phylogenetic tree construction pipeline. Ohana’s nemeco program estimates a rooted covariance matrix, where the root is arbitrarily chosen. Ohana’s convert program with cov2nwk option then recovers the full covariance matrix, computes the distance matrix, and approximates the distance matrix as a tree structure using the NJ algorithm. Finally, Ohana’s convert program with nwk2svg option renders the Newick tree in SVG format. For better control of the graphics, we recommend using our web service: http://www.jadecheng.com/graphs/

**Fig. 2.**
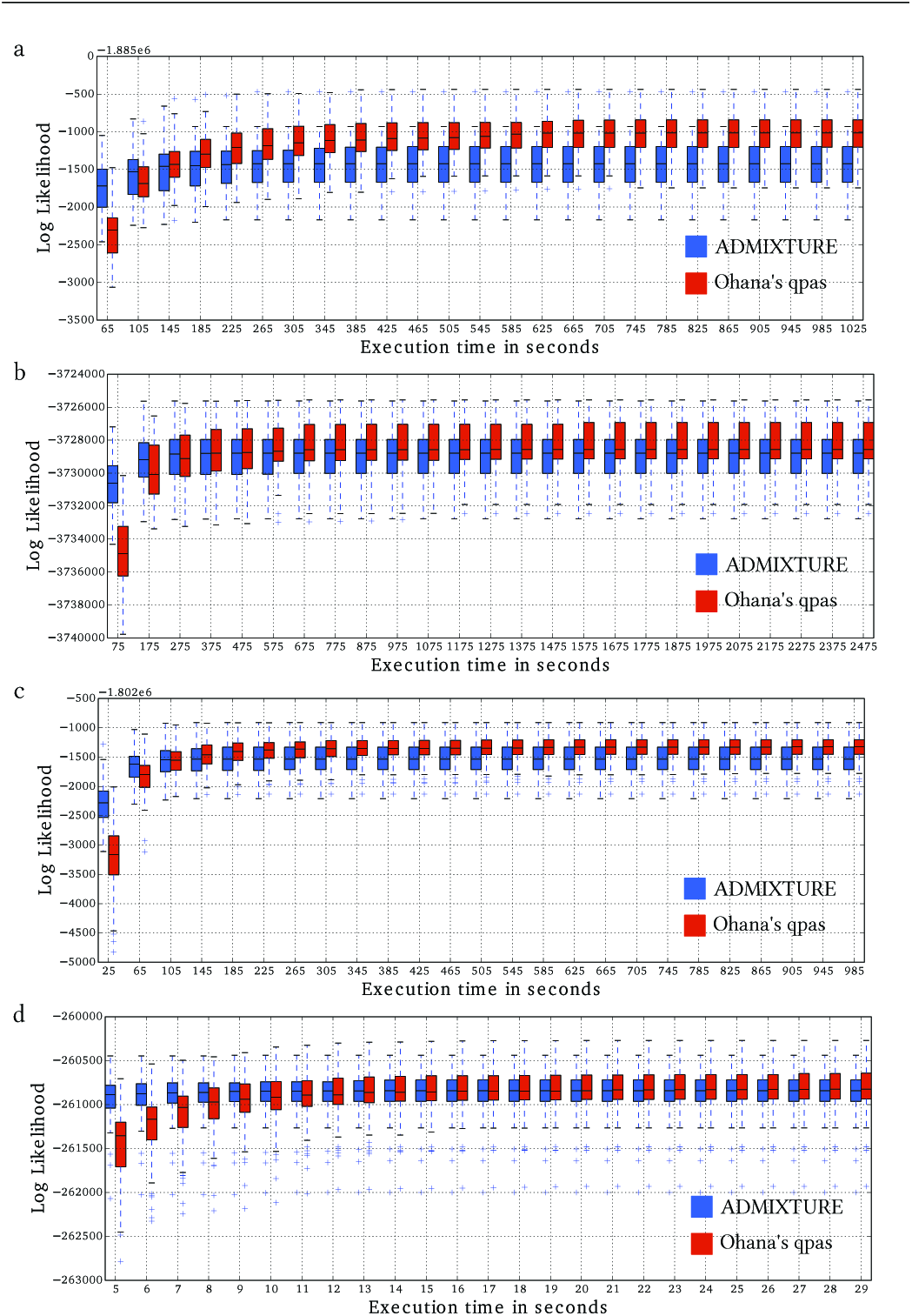
Comparison of computational speed and efficacy of ADMIXTURE and the qpas program in Ohana. The plots show the change in the distribution of log likelihood values, produced from the two programs over time. For each data set, each program was executed 100 times using random seeds (0, 1,…, 99) and *K* = 9. (a, b, c, d) are four different data sets, same as in Table 1.

### 3.2 Estimation of admixture fraction and tree on simulated data

We simulated data on a tree using coalescence simulations as described in the Methods section and estimated for different values of *K* (Figure 3). This mimics the procedure often used in real data analyses in which multiple values of *K* are explored and presented without knowing the true value of *K*, although this value can be estimated using a variety of methods (Alexander *et al.*, 2011; Scheet and Matthew, 2006;Wold, 1978).

**Fig. 3.**
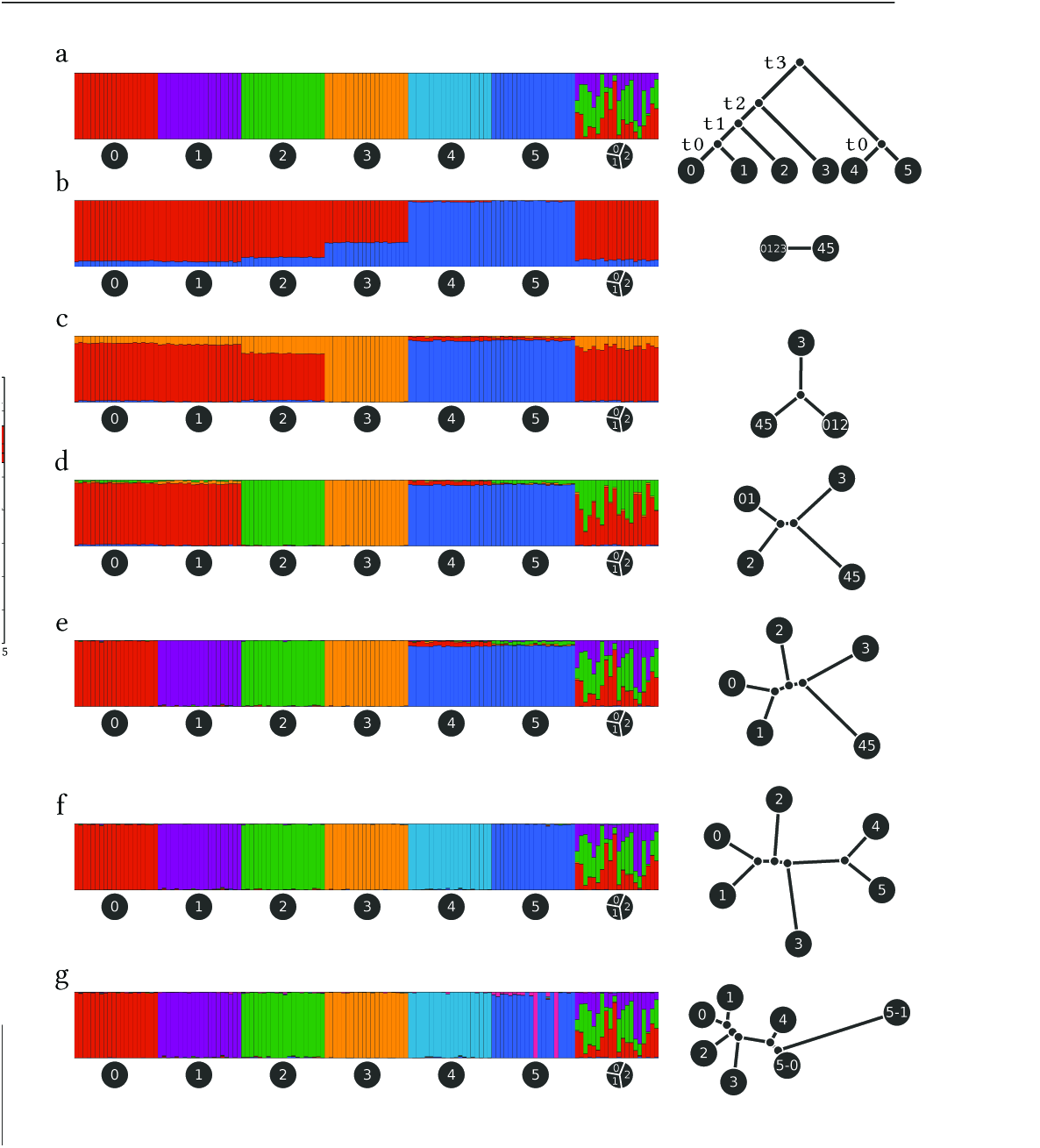
An evaluation of the tree inference procedure in Ohana using coalescence simulations. We simulated 140 individuals in 7 groups, 20 individuals per group. The first 6 groups were un-admixed. The last group was an equal mixture of the first 3 groups. (a) Simulated admixture (left) and simulated demography (right). (b, c, d, e, f, g) Estimated admixture (left) and estimated demography (right) for *K* = 2; 3; 4; 5; 6; 7, respectively. For each of the 6 populations, we simulated 100 sequences of size 20,000,000 bp using fastsimcoal2 (Excoffier *et al.*, 2013). We used a mutation rate of 2 × 10^˗8^ per generation, a recombination rate of 10^˗8^ per generation, and a population size of 50,000. The time parameters were 1000, 2000, 3000, and 4000 generations for *t*_0_, *t*_1_, *t*_2_, and *t*_3_, respectively. A total of 125,787 markers survived filtration for being polymorphic, diallelic, and with minor allele frequency greater than 5%. We then estimated admixture fractions and population trees using values of K ranging from 2 to 7.

The plots show good correspondence between the true and the estimated values, for both admixture proportions and demography. Furthermore, the changes in tree topology as *K* changes reflect the hierarchical structure of the tree. For example, at *K* = 4 the internal branch reflects the split between populations (0, 1, 2) and (3, 4, 5).

### 3.3 Model limitations

There are at least three reasons why tree estimation using a Gaussian model based on estimated allele frequencies may face challenges. First, the allele frequencies are treated as observed data, but they are truly estimates. This has the potential for introducing a variety of biases. Second, the use of a Brownian motion model to approximate genetic drift is inaccurate near the boundaries and for long divergence times, likely leading to underestimates of the lengths of long branches. Third, due to differences in sample sizes for different populations, the Structure model may not identify groups that correspond to natural units of a tree, even when the populations truly have evolved in a tree-like fashion.

We explore some of these issues in the following simulation study (Figure 4) by simulating trees with different divergence times: short, medium, and long. For very short divergence times (Figure 4-a), the covariance matrix was estimated poorly because of the small differences in allele frequencies across populations. This in turn leads to reduced accuracy in the estimation of the tree. While the topology is recovered correctly, the lengths of the external branches are overestimated. This likely happens because the Structure model tends to maximize allele frequency differences for finite sample sizes, i.e. the estimated difference in allele frequencies between pairs of populations tends to be larger than the true difference. This is an issue that can be mitigated with larger sample sizes and tends to be a problem only when branch lengths are very small. Nonetheless, it will likely affect many real data analyses.

**Fig. 4.**
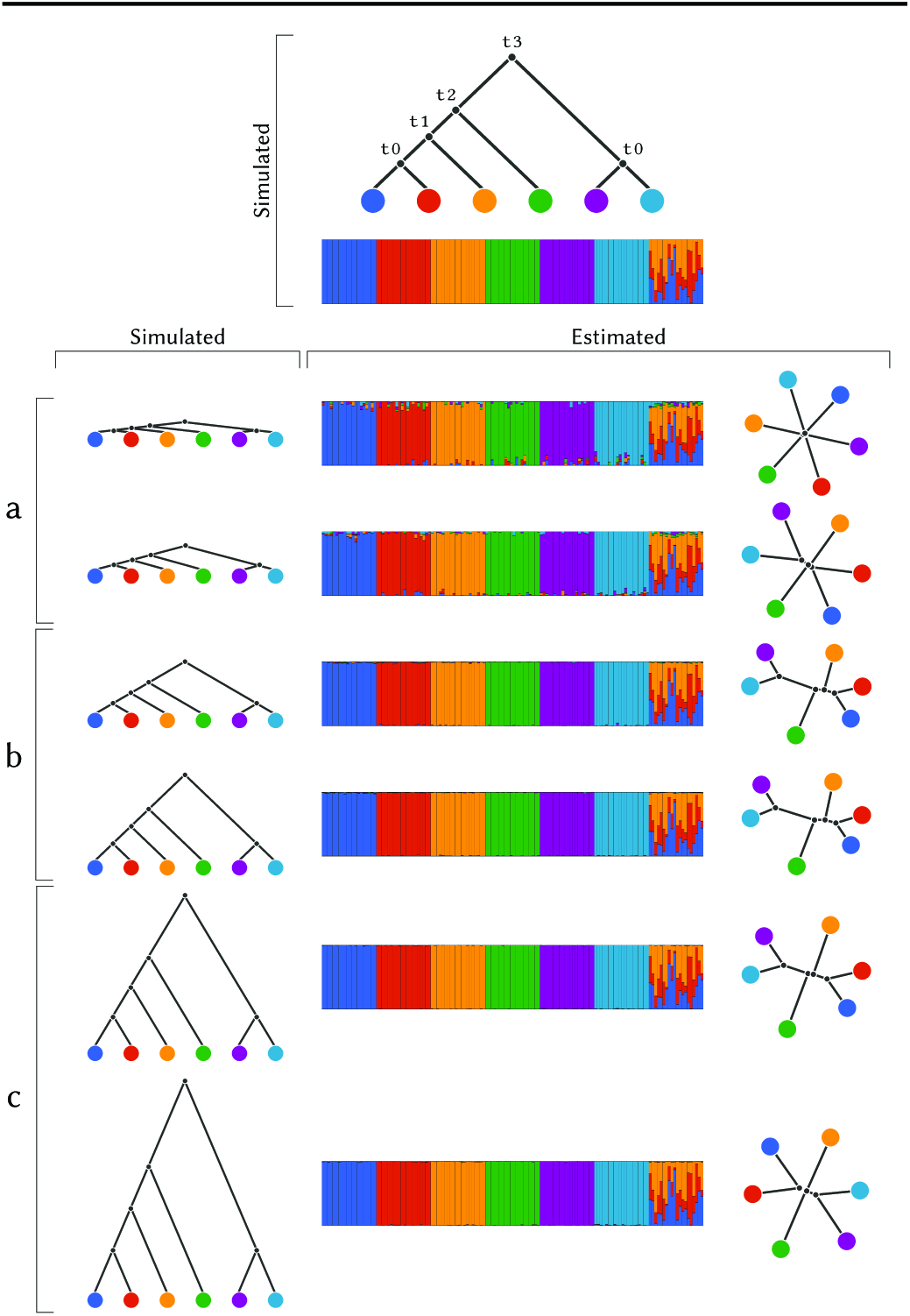
A simulation study for different divergence times. We simulated 140 individuals in 7 groups, 20 individuals per group. The first 6 groups were un-admixed. The last group was an equal mixture of the first 3 groups. We illustrate the simulated demography on the top. We simulated 6 divergence scenarios, 2 short shown in (a), 2 medium shown in (b), and 2 long shown in (c). From the shortest to the longest divergence scenario (top to bottom), the split times (*t*_0_, *t*_1_, *t*_2_, *t*_3_) in generation were: (10, 20, 30, 40), (100, 200, 300, 400), (1000, 2000, 3000, 4000), (1500, 3000, 4500, 6000), (10000, 20000, 30000, 40000), (20000, 40000, 60000, 80000).

In the long divergence scenario, Figure 4-c, another problem arises. For such long branches, the Brownian motion model is a poor approximation to genetic drift, and the mapping between the two transition probability functions (i.e. Wright-Fisher diffusion versus Brownian motion) is such that divergence times tend to be underestimated when they are long. The consequence is that the branch lengths of the tree are underestimated. We verify that this is the source of the bias by also simulating data under a Gaussian model directly and showing that under this model there is no significant bias for long branch lengths. This is described in SI Section 1. We note that the poor approximation of the Brownian motion model to the Wright-Fisher diffusion for long divergence times is a limitation for any inference system using similar statistical models such as TREEMIX (Pickrell *et al.*, 2012) and Bayenv (Gunther *et al.*, 2013), and it might be worthwhile in future work to explore the consequence of this effect for those methods as well.

In the medium-length divergence scenario (Figure 4-b), neither of the two previously mention sources of bias affect the inferences, and the estimates of the branch lengths are therefore quite close to the true values. In all three divergence scenarios, the tree topologies were always estimated accurately.

### 3.4 Other simulation scenarios

We also evaluated the performance of the method under several other simulation scenarios, and the results are presented in SI Section 2 to 5. A few noteworthy observations include: (1) In more than one simulation scenario with ancient admixture, the population was not inferred to be admixed but received a unique admixture component, SI Section 2 Figure 4 and Section 3 Figure 5. The probability of inferring admixture likely depends on the amount of drift since admixture. In the context of much human data showing evidence of ancient admixture, it might beworthwhile in future studies to explore how much drift after admixture is required to erase the signal of admixture. (2) When *K* is smaller than the true number of ancestry components, populations with few individuals represented in the sample tend to be (wrongly) inferred as admixed, SI Section 5 Figure 7. There is a clear dependence on sample size in inferences of admixture components in the Structure model. Similarly, the outgroup tends to be identified as the first admixture component that splits from the rest of the individuals, only when the outgroup is well-represented in the sample in terms of the number of individuals.

**Fig. 5.**
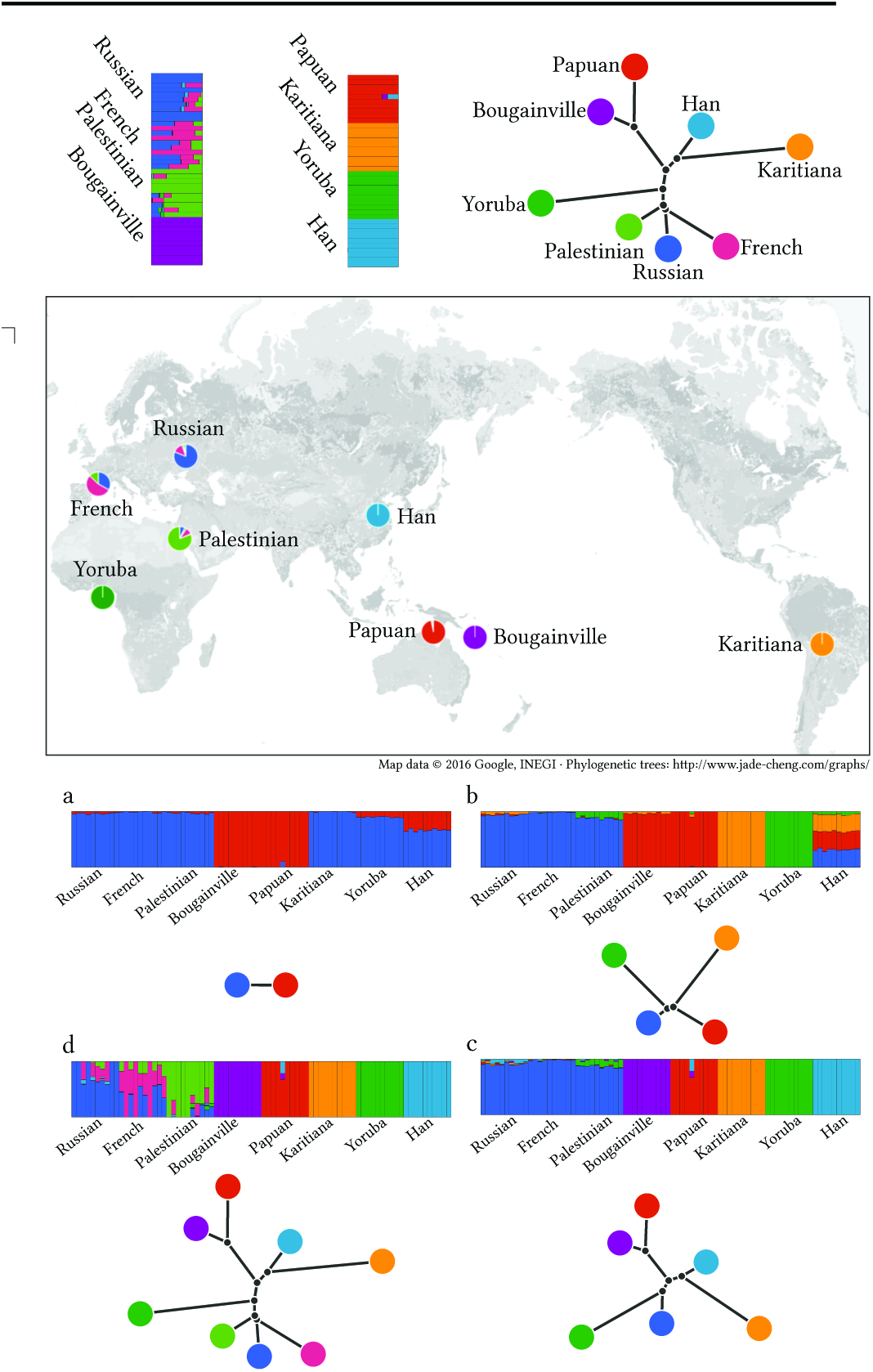
Analysis of human global data. We used a data set compiled from the HGDP project containing 80 individuals from 8 populations, 10 per population. We filtered markers using Plink (Purcell *et al.*, 2007) with options –indep 50 5 2 –geno 0.0 –maf 0.05. A total of 125,787 markers survived the filtration and were used for the analysis. For each *K* value, we dispatched 32 executions with random seeds from 0 to 31. We report only results from the execution that reached the best likelihood for each *K*. The plots show individual admixture proportions and population trees for several different values of K. The map combines the admixture results and geographical records of the HGDP samples. Each slice of each pie chart shows the sum of one component estimated in samples collected at that region. (a, b, c, and d) show the admixture and tree estimates for *K* = 2, 4, 6, 8, respectively.

### 3.5 Real data analysis

To illustrate the method, we apply it to the panel of global human data described in the Methods section (Figure 5), using a range of *K* values. The topologies of the trees largely mimic what is already known about human ancestry (e.g., (Reich *et al.*, 2012)), i.e. using a root in Africa, Asians and Native Americans cluster together, the European and middle Eastern groups cluster together, etc. In addition to Yorubans having a long branch because this group is an outgroup to the rest, we also notice a relatively long branch leading to Native Americans, reflecting the increased drift in this group due to the bottleneck into the Americas and possibly small population sizes thereafter.

## 4 Discussion

In this paper, we introduced a new implementation of the Structure model in a maximum likelihood framework. We compared the new optimization algorithm to the one implemented in the hitherto fastest program, ADMIXTURE. The qpas program in our software, Ohana, generally outperformed ADMIXTURE by obtaining estimates with higher likelihood values in similar computational time.

In addition, we presented a new approach for estimating trees for ancestry components. Using coalescence simulations, we showed that when the trees are interpreted as reflecting true population trees, external branch lengths tend to be overestimated for small divergence times. However, for long divergence times, the use of a Gaussian model and its inaccuracy in approximating genetic drift cause branch length estimates to be downward biased. Nonetheless, the estimates of tree topology appear reasonably robust. The tree estimation and visualization tool should be of use to other researchers as an additional possible component of a Structure model analysis of the data. The tree is a visualization of the covariance structure of the admixture components, and it may as such be useful even if a strict interpretation of a evolutionary tree may not be warranted. There might be several reason why such an interpretation may not be appropriate, most of all because the true nature of the evolution of the ancestry components may not be well-described by a tree. Ancestry components are constructions that may or may not reflect true ancestral populations.

## Acknowledgements

Thiswork is funded by the Danish Council of Independent Research Sapere Aude grant 12-125062; Conflict of Interest: none declared.

